# The Relationship between Institutional Prestige, Journal Impact Factor, and Sample Size in fMRI Studies of Memory: A Systematic Review

**DOI:** 10.64898/2026.04.24.720672

**Authors:** Moe Mansour, Samuel P. Chipman, Ariana M. Hedges-Muncy, Nathan M. Muncy, C. Brock Kirwan

## Abstract

Low statistical power remains a persistent concern in functional magnetic resonance imaging (fMRI) research, largely due to small sample sizes. Although prior work has documented gradual increases in sample size over time, it remains unclear whether structural factors in the publication process are associated with study design characteristics such as sample size. This review addresses this gap by analyzing a large sample of fMRI studies to assess how institutional prestige, journal impact factor, and journal review practices are associated with sample size. We analyzed articles published in 2021-2024 reporting new fMRI data collection in adult humans and including a measure of memory. We found studies with specialized populations, such as patient populations, had smaller sample sizes, as did studies with task-based designs compared to resting-state designs. We also found larger sample sizes were associated with journals with a double-blind review process. Institutional prestige was positively associated with sample size such that more highly ranked institutions tended to have larger samples, but there was no interaction between review type (single-vs. double-blind) and prestige, indicating this difference is not likely due to reviewer bias. Journal impact factor was not associated with sample size, however institutional prestige score predicted journal impact factor. These results suggest structural factors at the institutional level likely have a stronger influence on published study sample size than reviewer practices or biases.

---

Cognitive neuroscience studies suffer from low sample sizes, particularly those employing functional magnetic resonance imaging (fMRI) techniques (Button et al., 2013; Simmons et al., 2011). Studies with smaller sample sizes tend to report more activation foci than meta-analyses that combine data across studies (David et al., 2013), possibly reflecting greater false discoveries with smaller sample sizes (Button et al., 2013). Recent empirical studies have demonstrated that small sample sizes reduce the likelihood of observing brain-behavior correlations (Grady et al., 2021) and reduce the replicability of task-based fMRI designs (Turner et al., 2018), which may be compounded for some analysis pathways. For example, Bossier and colleagues (2020) examined activation maps for subsets of participants drawn from a larger cohort study and demonstrated poor test-retest reliability when comparing thresholded activation maps but more promising results when comparing correlations between un-thresholded maps. Similarly, Kampa and colleagues (2020) showed high replicability when considering activations at the whole-brain level (particularly for tasks with strong activation) but not at the level of individual regions of interest.

The fields of psychology and cognitive neuroscience have responded to the replication crisis by increasing sample size over time. Within psychology, median sample sizes have increased from 40 in 1995 to 120 in 2019 (Bakker et al., 2025). Specific to the fMRI literature, Carp (2012) reported a median sample size of 15 for a random sample of 300 papers published between 2007 and 2012. In a survey of fMRI studies from 2017, Yeung reports a median sample size of 33 (Yeung, 2018). Szűcs and Ioannidis (2020) demonstrated sample sizes in fMRI studies have increased at a rate of about 0.74 participants per year between 1993 and 2018.

Here we asked whether markers of prestige such as authors’ institution ranking or journal impact factor and journal review practices are related to sample size of fMRI studies examining long-term memory (our particular research area). Prior research has shown the prestige of a researcher’s training institution is a strong predictor of subsequent career placement, often outweighing individual productivity measures (Pinheiro et al., 2017). More broadly, prestige functions as a form of symbolic and social capital within academia, influencing how researchers are perceived and evaluated within hierarchical networks (Zeitlyn & Hook, 2019). As a result, affiliation with more prestigious institutions is associated with advantages in academic placement (Pinheiro et al., 2017) and may influence how research is evaluated within academic networks (Zeitlyn & Hook, 2019), suggesting that prestige may influence reviewers’ evaluations when they have access to that information. Similarly, journal impact factor is widely used as an indicator of a journal’s visibility and influence.

Despite growing attention to sample size and replicability in cognitive neuroscience, it remains unclear whether structural factors in the publication process are associated with study design characteristics such as sample size. This review addresses this gap by analyzing a large sample of fMRI studies to assess whether institutional prestige and journal impact factor are associated with sample size. We hypothesized when reviewers have access to authors’ institutional affiliation, as in the case of single-blind review, institutional prestige would be negatively associated with sample size. Conversely, we hypothesized when the review process was double-blinded, there would be no relationship between authors’ institutional prestige and sample size. Further, we hypothesized studies with larger sample sizes would be more likely to appear in journals with higher impact factor scores.

## Methods

### Data Selection

We identified articles in the PubMed database with the words “FMRI” and “memory” in the title or abstract and the terms “adult” and “humans” in the Medical Subject Headings [MeSH] field for papers published between 2021 and 2024 according to the PubMed entry date [edat] field. The final search for records was performed on November 3, 2025 and resulted in 565 total articles, which were all retrieved for further evaluation. Articles were included if they were primary research studies published as peer-reviewed articles in English that reported collecting both fMRI data and a behavioral measure of long-term memory. Studies were excluded if they reported analyses of previously collected datasets (e.g., the Human Connectome Project), if they included data from children or animal models, or if they were protocol papers, meta-analyses, or reviews. Initial screening for inclusion/exclusion criteria was performed by an LLM (Microsoft Copilot) and was spot-checked by the authors, who randomly tested papers to verify the inclusion/exclusion reasoning and determination. After exclusions, the final dataset included 338 articles. Figure 1 depicts the PRISMA 2020 article selection process (Page et al., 2021).

**Figure 1.**
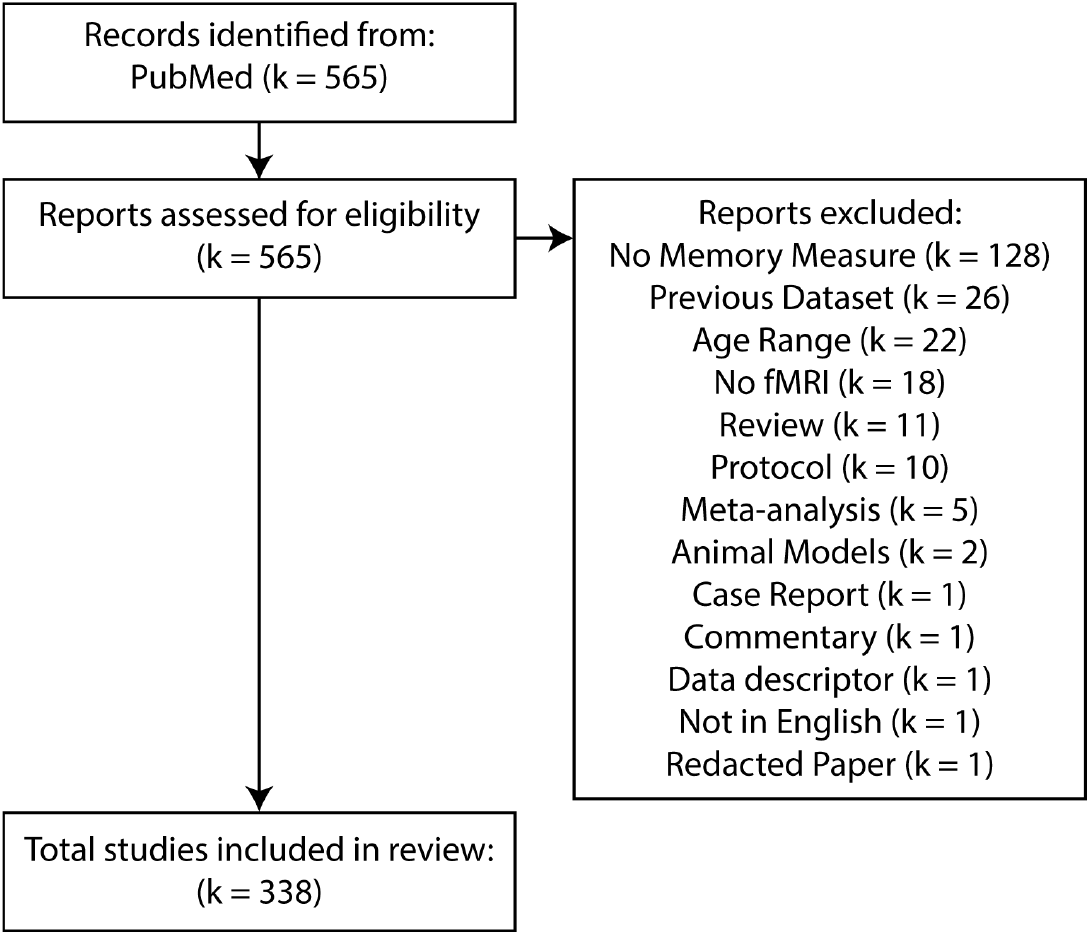
PRISMA 2020 flow diagram of study identification and assessment for inclusion.

We processed all included articles for first and last author names, h-index values, and institutional affiliations, as well as journal impact factors and review process. First and last author names and affiliations were manually extracted. If there was more than one equal contributor, we used the first and last authors listed, respectively. Similarly, when multiple affiliations were listed, we used the first affiliation listing. Institution ranking was taken from the 2025 U.S. News Best Global Universities rankings (https://www.usnews.com/education/best-global-universities/). Institutions without rankings were entered as N/A. H-index values were retrieved from Scopus using current (i.e., 2025) values. Journal impact factors were taken from the 2024 Journal Citation Reports (https://clarivate.com/). Journal review type (single-vs. double-blind) was determined from individual journal submission websites.

Each included article was analyzed for final sample size, population type (patient vs. control), number of groups, fMRI paradigm (resting-state vs. task-based), the number of MRI scan sessions, and the number of distinct experiments using an LLM (Microsoft Copilot). Extracted values were manually verified through spot-checking (approximately 10%) of selected articles to confirm accuracy of extracted values. When discrepancies or ambiguities were identified (2.1% of articles), values were resolved through manual review of the full-text article.

### Data Cleaning and Organizing

We conducted all data processing and analyses using R (4.5.1) in Jupyter Notebooks. Institutional rankings are traditionally coded as lower numbers indicating better rankings. Accordingly, we reverse-coded rankings such that higher numbers were now associated with better rankings. We next computed missingness in the data and found 15.1% and 13.6% of first and last author institution rankings were missing, respectively. H-index values had 0.6% missingness, but no other variables had any missing values. Missing values were nonparametrically imputed through a random forest model. The model had an out-of-bag error of 17.9%. Finally, we adjusted each study’s final sample size by the number of unique scanning session, number of groups, and the number of experiments included in the article. That is, the adjusted sample size *y*^∗^ is

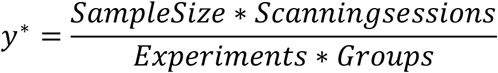

Adjusted sample sizes were winsorized at the 1st and 99th percentiles to account for extremely large and extremely small sample sizes. Values outside this range were replaced with the respective percentile cutoffs, preserving all observations while limiting the impact of outliers.

### Data Analysis

As first and last author institution rankings and h-index values were assumed to be strongly correlated, we performed a principal component analysis (PCA) to account for multicollinearity and capture shared variance across these related metrics. The first PCA component accounted for 43.4% of the variance. Institutional rankings had strong negative loadings on this component (both -0.67) while author h-index values had weaker negative loadings (-0.09 and -0.30). To account for these negative loading, we inverted this factor to create a more intuitive Prestige score where higher values reflected higher institutional prestige and author citation count.

To examine the association between prestige-related measures and adjusted sample size, we fit a gamma regression with a log link using adjusted sample size as the dependent variable and using prestige score, population type (patient vs. control), journal review type (single-vs. double-blind), fMRI paradigm (resting-state vs. task-based), journal impact factor, and the interaction of review type by prestige score as independent variables. Finally, we fit a second gamma regression with a log-link using journal impact factor as the dependent variable and prestige score, population type, review type, and fMRI paradigm as independent variables.

### Data Availability

All data and analysis code are available here: https://osf.io/6txuh/overview?view_only=7bcb9e0fe5fe41f29015fae4f514a54c.

## Results

The distribution of institution rankings, author h-index values, and journal impact factors along with adjusted sample size are depicted in Figure 2. Median sample size was 32 (1st quartile = 21.6, 3rd quartile = 59.6). As predicted, there were significant correlations between first and last authors’ institution rankings (r = 0.67, t(336) = 16.37, p < 0.001), and h-index values (r = 0.17, t(336) = 3.09, p < 0.01), supporting the decision to create prestige factors using PCA for the main analysis.

**Figure 2.**
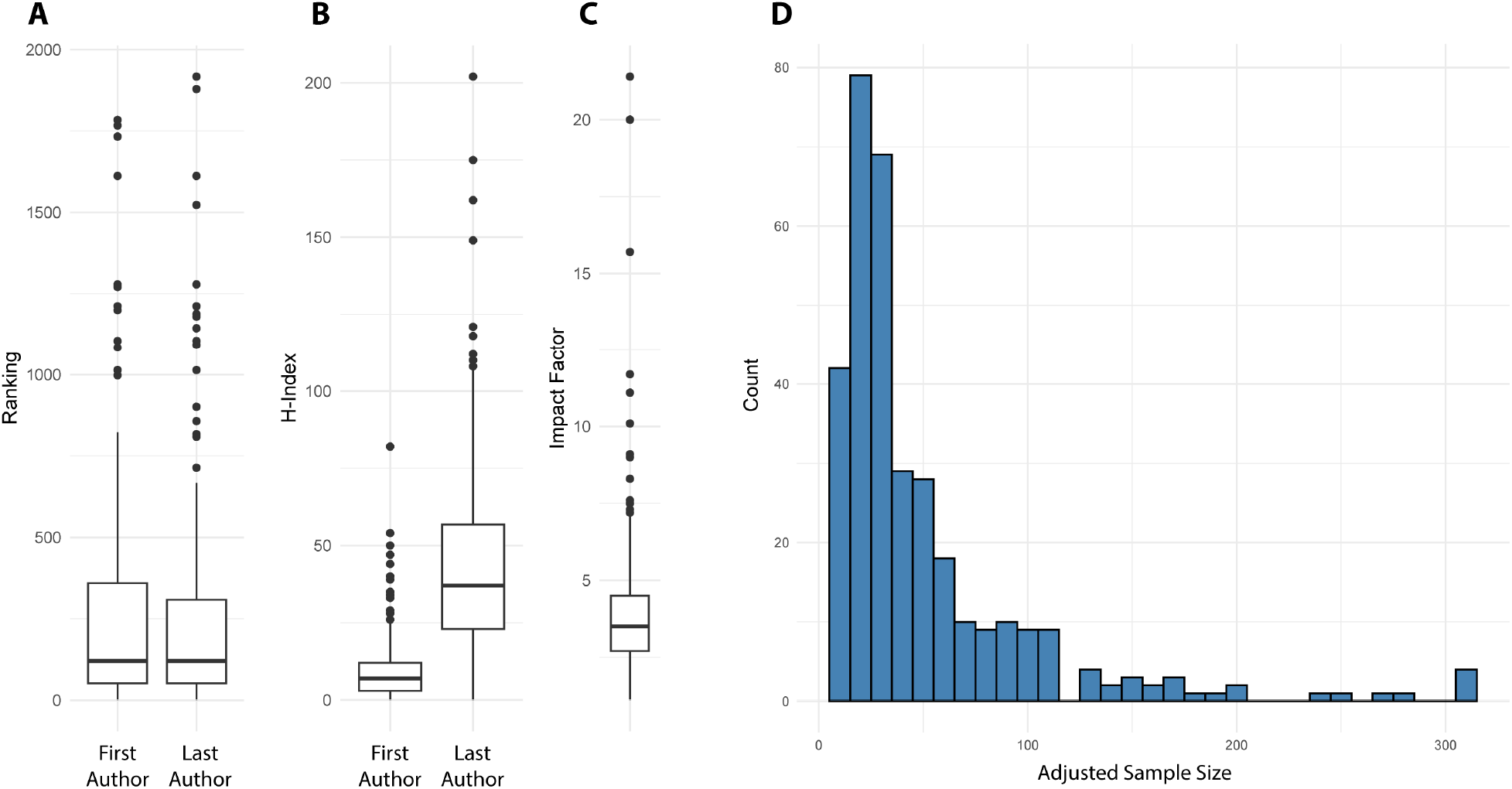
Final sample characteristics for 338 primary research articles examining long-term memory published between 2021 and 2024. A) Distribution of institutional rankings and B) h-index values for first and last authors. C) Journal impact factors. D) Histogram of adjusted fMRI sample sizes

Table 1 reports the results of the GLM model predicting sample size. Prestige score was a significant positive predictor of sample size (p < 0.05), indicating that authors associated with higher prestige institutions tend to have publications with larger sample sizes (Figure 3). Review type was a negative predictor of sample size, indicating that journals with double-blind review tend to publish studies with larger sample sizes. Study population was also a significant predictor of sample size, with studies focused on patient or other specialized populations having smaller sample sizes. Task paradigm was also a significant predictor, with task-based studies having smaller sample sizes than resting-state studies. Journal impact factor was not a significant predictor of sample size. Finally, the interaction of review type by prestige was not significant, indicating that reviewers’ awareness of authors’ identity or institution does not affect the sample size of published studies.

**Table 1.**
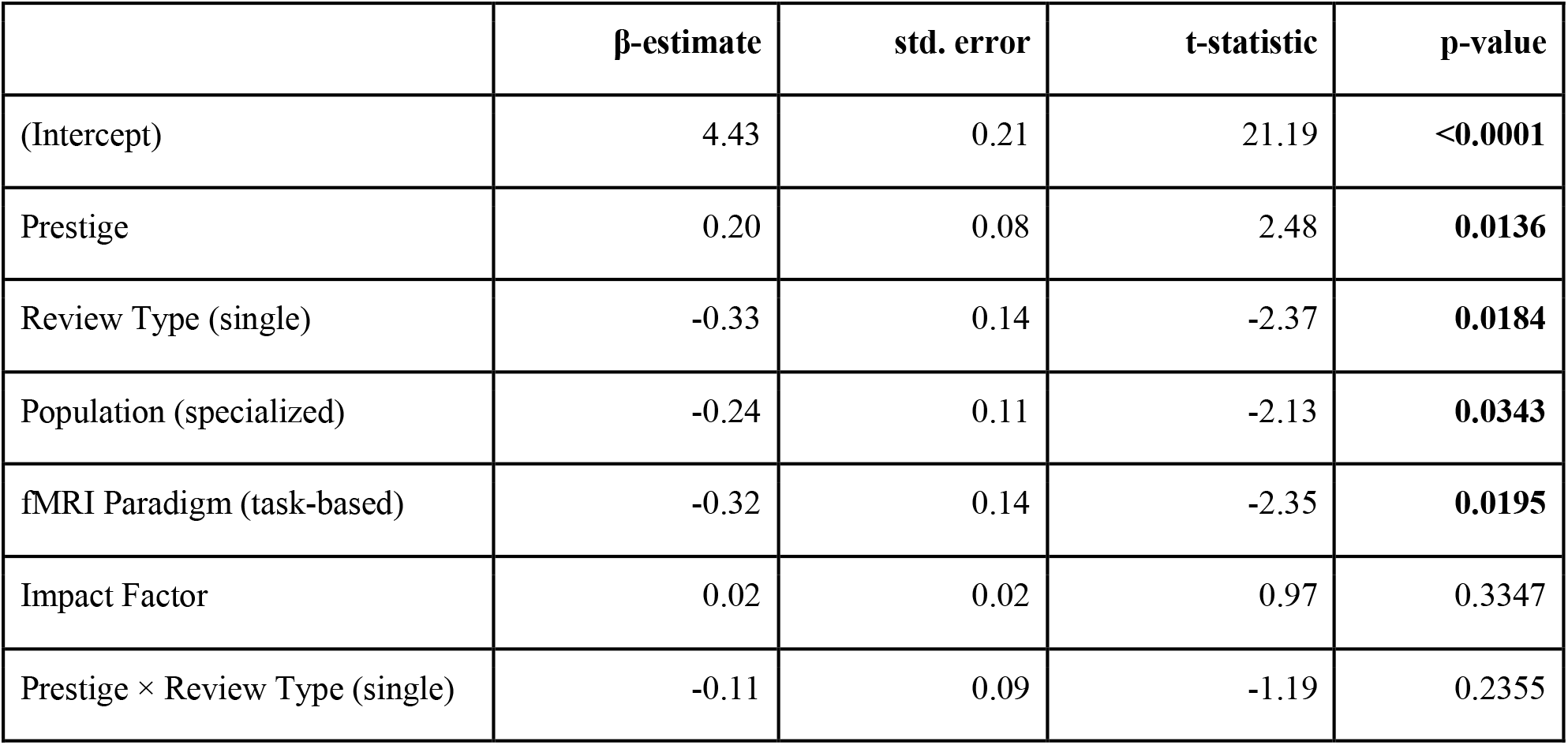
Parameters of the model predicting sample size.

**Figure 3.**
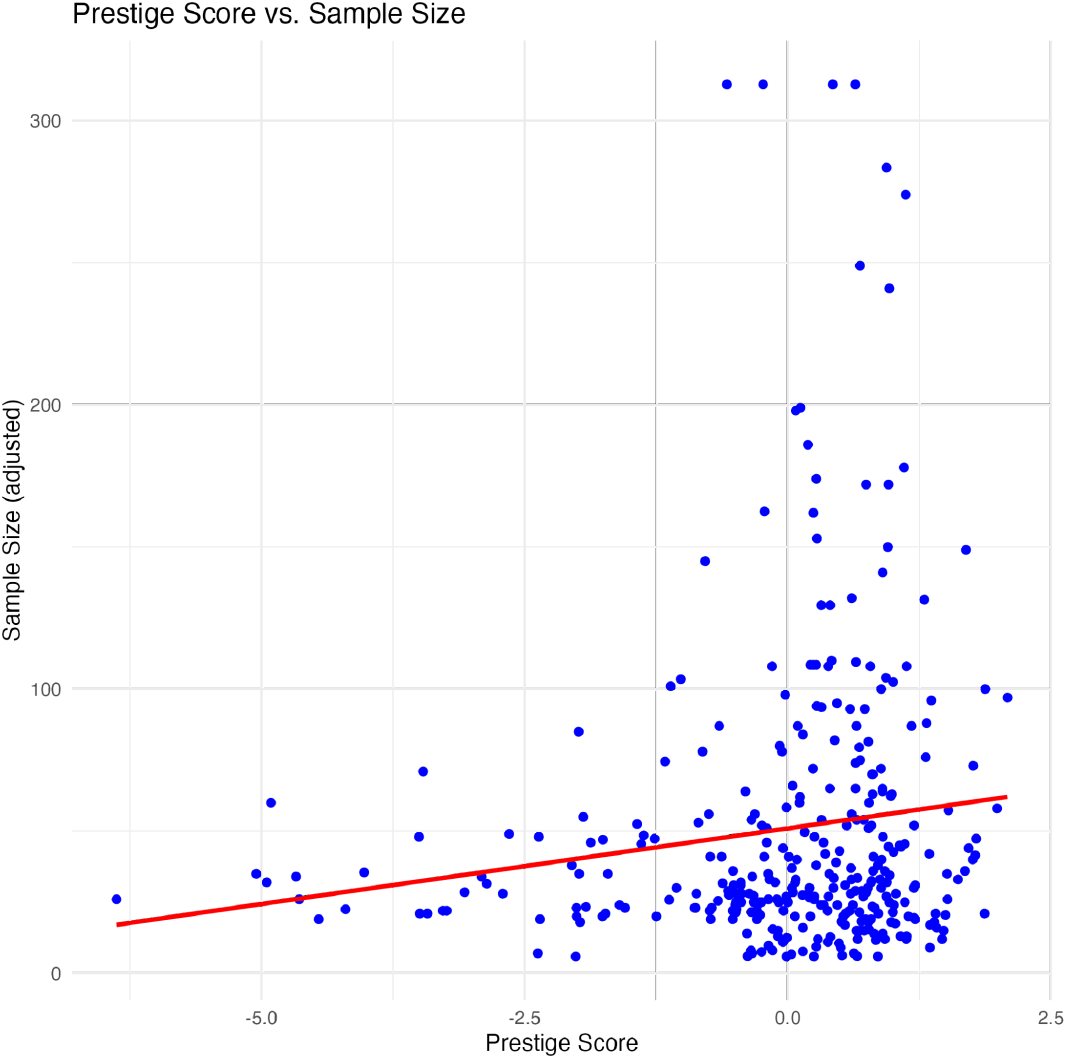
A composite prestige score derived from first and last author institution ranking and h-index values positively predicted adjusted sample size in fMRI studies.

Table 2 reports the results of the GLM model predicting journal impact factor of the studies in our dataset. Prestige was a significant predictor of journal impact factor (p < 0.001), indicating that authors from higher prestige institutions tend to publish in journals with higher impact factor scores (Figure 4). Review type was also a significant predictor of impact factor. Study population and fMRI paradigm did not predict journal impact factor. Finally, there was a significant interaction between review type and prestige but with a negative coefficient, indicating prestige was more predictive of impact factor in double-blind journals than in single-blind journals.

**Table 2.** Parameters of the model predicting impact factor.

| | β-estimate | std. error | t-statistic | p-value |
| --- | --- | --- | --- | --- |
| (Intercept) | 1.76 | 0.12 | 15.26 | < 0.001 |
| Prestige | 0.24 | 0.05 | 4.79 | < 0.001 |
| Review Type (single) | -0.44 | 0.08 | -5.22 | < 0.001 |
| Population (specialized) | 0.01 | 0.07 | 0.19 | 0.8478 |
| fMRI paradigm (task) | 0.05 | 0.09 | 0.60 | 0.5503 |
| Prestige × Review Type (single) | -0.17 | 0.06 | -2.87 | 0.0044 |

**Figure 4.**
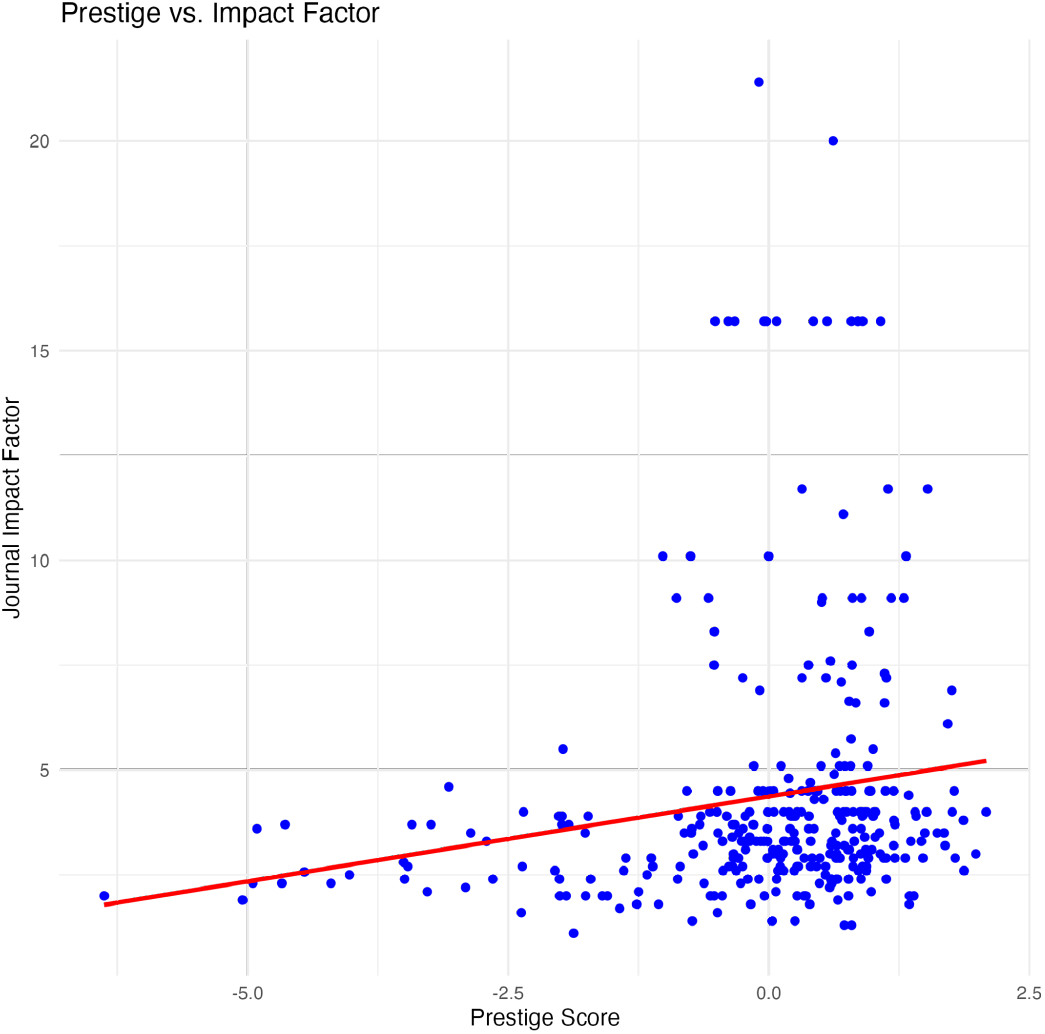
Prestige score was a significant predictor of journal impact factor, with studies from higher prestige institutions more likely to be published in high impact factor journals.

## Discussion

We examined the factors associated with sample size of fMRI memory studies published in the recent neuroscience literature. Prestige was a consistent predictor of study sample size, with authors from higher prestige institutions publishing studies with larger sample sizes. Other factors influencing sample size included the study population (with larger sample sizes associated with control participants than patient populations), fMRI paradigm (with larger sample sizes associated with studies that employed only resting-state designs), and review type (with larger sample sizes associated with double-blind reviews). However, there was not a review type by prestige score interaction, indicating reviewers’ knowledge of the authors’ institution did not influence their evaluation of study quality (as indexed by study sample size), contrary to our original hypothesis. The journal impact factor of fMRI memory studies was also significantly predicted by institutional prestige and by review type (and their interaction) but not by fMRI paradigm or population type.

We originally hypothesized markers of prestige such as institutional ranking or author h-index would have an influence on the sample size of published results, particularly when reviewers were aware of these markers in single-blind reviews. Our results do not support this hypothesis; indeed, the relationship between prestige score and sample size was the opposite of that hypothesis. The lack of interaction between review type and prestige on study sample size indicates reviewers do not consider markers of prestige when evaluating study quality (at least as indexed by sample size). The positive relationship between prestige and sample size might suggest institutional resources or research infrastructure associated with prestige may facilitate larger studies.

Other than prestige score, factors that influence fMRI study sample size included study population and experimental design. Studies that included specialized populations (such as patient populations) were smaller than those that only included healthy control participants. Similarly, studies employing a task-based design tended to have smaller sample sizes than those using a resting-state-only design. In both cases, ease of recruiting and ease of data collection likely contribute to final sample size. In contrast, journal impact factor was not a significant predictor of adjusted sample size when controlling for prestige, population type, and fMRI paradigm. Together, these results may indicate journal impact factor may not be a strong indicator of the strength of study design, but rather journal impact factor may reflect other characteristics of publication venue.

Higher institutional prestige was associated with publications in journals with higher impact factors. Interestingly, this association was stronger in journals with double-blind review than in those with single-blind review. Thus, it seems unlikely reviewers’ knowledge of institutional ranking influences their recommendations regarding acceptance decisions. Rather, a more likely explanation is authors associated with higher prestige institutions tend to submit to journals with higher impact factors, which also tend to employ a double-blind review process. It should be noted that even when journals employ double-blind review, the initial editorial assessment of whether to send a manuscript out for review is not blinded for author identity, so it cannot be ruled out from these data that prestige has an effect on editorial review decisions.

Some limitations to the present study should be noted. First, journal impact factor is not the only measure of a journal’s influence or reputation. Other metrics include Scimago Journal Rank (SJR), which ranks journals weighted by the reputation of citing journals (González-Pereira et al., 2010), and Eigenfactor score (Bergstrom, 2007; Bergstrom et al., 2008). Some have noted IF is influenced by a few highly cited papers in a journal (e.g., Seglen, 1997). Given these caveats, it is still notable that IF is positively correlated with other measures of journal impact and reputation (Chang et al., 2011; Ward, 2014). Also, sample size is not a perfect proxy for statistical power in fMRI studies. Nee (2019) demonstrated task-based fMRI activations are replicable with modest sample sizes (n=16) as long as sufficient data are collected within subjects (Chen et al., 2022). We did not evaluate studies in the current sample for the number of trials or runs obtained within subjects.

Additionally, our sample may be biased by several factors as evidenced by the presence of few “low” ranked institutions in our sample. Higher prestige institutions are more likely to have research-dedicated MRI scanners, which may limit the number of fMRI papers submitted by authors at lower prestige institutions. This access difference has been partially addressed through recent advancements in open-science practices which have made large cohort datasets available to the broader research community. However, as we chose to focus on newly published studies, we excluded papers that analyzed data from these sources. Finally, although PubMed offers extensive coverage of the biomedical research literature, there are omissions in the database (Frandsen et al., 2025). Further studies may wish to examine whether there are systematic differences between studies indexed in PubMed compared to those that are not.

## Conclusion

This review examined whether prestige and journal impact factor are associated with sample size in adult fMRI studies of memory. Overall, prestige-related measures were a significant predictor of sample size, with higher prestige institutions associated with larger sample size. Journal review type, study population, and fMRI paradigm were also significant predictors of sample size. Journal impact factor was not a significant predictor of sample size, but prestige-related measures were in turn significant predictors of journal impact factor. Together these results indicate factors associated with prestige such as institutional research resources are likely better predictors of study sample size than journalistic review processes.

